# Benchmarking interface confidence as a selection criterion for computational nanobody design against survivin’s BIR-domain groove

**DOI:** 10.64898/2026.10.04.753929

**Authors:** Charlotte Gately, Xinyao Guan, Xianting Ding

## Abstract

Designed binders are routinely filtered on predicted interface confidence, which reports how self-consistently a model places an interface, not where it lies. We designed 200 nanobody loops against the survivin BIR-domain groove, co-folded 196, and checked epitope contact in every predicted complex. Confidence showed no detectable association with epitope engagement (ρ = +0.100, 95% CI −0.04 to +0.24, *n* = 196); its highest and lowest quartiles failed alike (odds ratio 0.97, 95% CI 0.41 to 2.26). It tracked instead the run’s other confidence estimates (ρ = +0.61). Twelve-seed replication separated two candidates the criteria had not: one reached the complete epitope in 10 of 12 runs, the other never. Across 155 predictions the top-ranked model was no better than one drawn at random from its five (*P* = 0.610), though only 3 of 31 failed on every model. Nothing was assayed; nb_cand_057 is an in silico candidate only.

## 1. Introduction

Survivin, encoded by *BIRC5*, is the smallest inhibitor of apoptosis protein: 142 residues, a single baculovirus IAP repeat (BIR) domain and a C-terminal helical extension mediating homodimerisation (Verdecia et al., 2000; Chantalat et al., 2000). It both directs chromosome alignment and cytokinesis in the chromosomal passenger complex (Wheatley and Altieri, 2019) and opposes apoptosis by binding Smac/DIABLO, preventing it from derepressing caspase activity (Song, Yao and Wu, 2003).

Its expression profile is what makes it a target. Survivin is abundant in most human malignancies and greatly reduced in differentiated adult tissue (Ambrosini, Adida and Altieri, 1997), though expression persists in proliferating normal compartments (Fukuda and Pelus, 2006), so any therapeutic window rests on degree rather than absence. Overexpression correlates with poor prognosis and treatment resistance across tumour types (Altieri, 2008; Wang and Greene, 2024), yet two decades of effort have produced no approved agent. The obstacles follow from the protein: it has no enzymatic active site, both functions run through protein interactions rather than catalysis, and both are discharged inside the cell.

The BIR domain, spanning approximately residues 15 to 89, presents a surface groove that engages short peptides by recognising a free N-terminal alpha-amino group (Du et al., 2012). Two physiological ligands use it: the AVPI tetrapeptide of mature Smac/DIABLO, and the T3-phosphorylated N-terminal tail of histone H3, which recruits survivin to centromeric chromatin (Jeyaprakash et al., 2011; Du et al., 2012). Du et al. (2012) determined crystal structures of both complexes, PDB 3UIH and 3UIG, showing that both engage the same pocket through the same chemistry, with binding preference reversing under defined point mutations within it. The groove is therefore structurally characterised and demonstrably tunable, and an exogenous binder occupying it intervenes at the binding step itself rather than upstream of it.

One residue in this surface carries functional validation obtained independently of crystallography. Song, Yao and Wu (2003) showed that the single substitution D71R abolishes the survivin-Smac/DIABLO interaction and eliminates survivin’s anti-apoptotic activity in taxol-treated cells. A computationally derived contact list is a hypothesis about which residues matter. Asp71 is the one residue in this epitope for which that hypothesis has already been tested experimentally, so we treat it throughout as a functional criterion rather than one contact among eleven.

Wang and Greene (2024) review the therapeutic strategies directed at survivin. Transcript-level approaches, small molecules acting through associated pathways and immunotherapies have all been pursued without an approved agent. Peptide agents are the most relevant category: Pathania et al. (2026) identified a previously unrecognised Smac-binding sequence within survivin and built a 25-residue survivin-derived peptide which, in stabilised cell-penetrating form, disrupts survivin dimerisation across multiple cancer cell lines. Two things follow for the present work: the interaction surface is engageable by a designed macromolecule, and intracellular delivery is achievable. No antibody-format binder against a survivin surface appears anywhere in that review.

Antibodies against survivin exist and are widely used, but binding survivin somewhere is not the same as binding this surface. Searching the published literature, we found none epitope-mapped to the BIR-domain groove, and none reported as an inhibitor of the survivin-Smac/DIABLO interaction in cells. The availability of detection reagents is readily mistaken for the availability of an inhibitor.

Diffusion-based generative models can now design binders to specified epitopes without a natural template. RFdiffusion (Watson et al., 2023) introduced hotspot conditioning, in which the designer nominates the target residues a binder must contact. Bennett et al. (2026) extended the approach to antibodies, holding the humanised camelid scaffold h-NbBcII10FGLA (Vincke et al., 2009) fixed and producing VHH and scFv binders with atomically accurate predicted interfaces. We adapt the logic of that approach rather than its machinery. Fixing the framework and varying one loop reduces the problem to a size that can be explored exhaustively on free computing resources, and means every candidate shares an identical scaffold, so interface contributed by non-designed regions can be separated from the contribution of the design.

Scoring is where such pipelines depend most heavily on structure prediction. AlphaFold2 (Jumper et al., 2021), its multimer extension (Evans et al., 2021) and the ColabFold implementation (Mirdita et al., 2022) co-fold candidate and target, and the resulting interface confidence estimate, ipTM, then filters which designs proceed to synthesis and testing (Watson et al., 2023; Bennett et al., 2026; Overath et al., 2025). ipTM reports how certain the model is that it has predicted an interface correctly. It does not report where on the target that interface lies, and it is not an estimate of affinity. A prediction can therefore be confident and wrong in the way that matters, placing a well-packed interface on the wrong face of the target.

Empirical grounds for caution have recently become available. Overath et al. (2025) assembled 3,766 experimentally characterised binders across fifteen structurally diverse targets, re-predicted every complex under AlphaFold2, AlphaFold3 and Boltz-1, and extracted more than 200 features per design. They found that ipTM is outperformed by interface-focused alternatives, notably the AlphaFold3-derived ipSAE, and that predictive performance varies substantially by target. They open by noting that prioritisation of designed binders still relies on heuristics. Their study asks whether a design binds at all, not whether it binds in the intended place, and aggregating fifteen heterogeneous targets cannot resolve the relationship for any one of them.

Two gaps motivate this work. The relationship between interface confidence and epitope-specific engagement has never been quantified across a complete design library against a single target, although it is answerable from output such a pipeline already produces and although the choice of filter decides which designs are ever made. And no antibody-fragment binder has been designed against the survivin BIR-domain groove or epitope-mapped to it, despite the surface being crystallographically defined and validated by mutagenesis, which makes it a target where engagement of the intended site can be checked residue by residue rather than assumed.

We therefore designed and evaluated a complete 200-member CDR3 library on the h-NbBcII10FGLA framework, verifying predicted epitope contact in every co-folded complex, and used the result to measure how far interface confidence, and a single prediction, can be relied on to identify a candidate that engages the intended site. This is a benchmark of selection criteria rather than a therapeutic design campaign. Nothing reported here was expressed or assayed, every candidate discussed is a computational candidate, and the design work serves the measurement rather than the other way round. Figure 1 sets out the pipeline in full.

**Figure 1.**
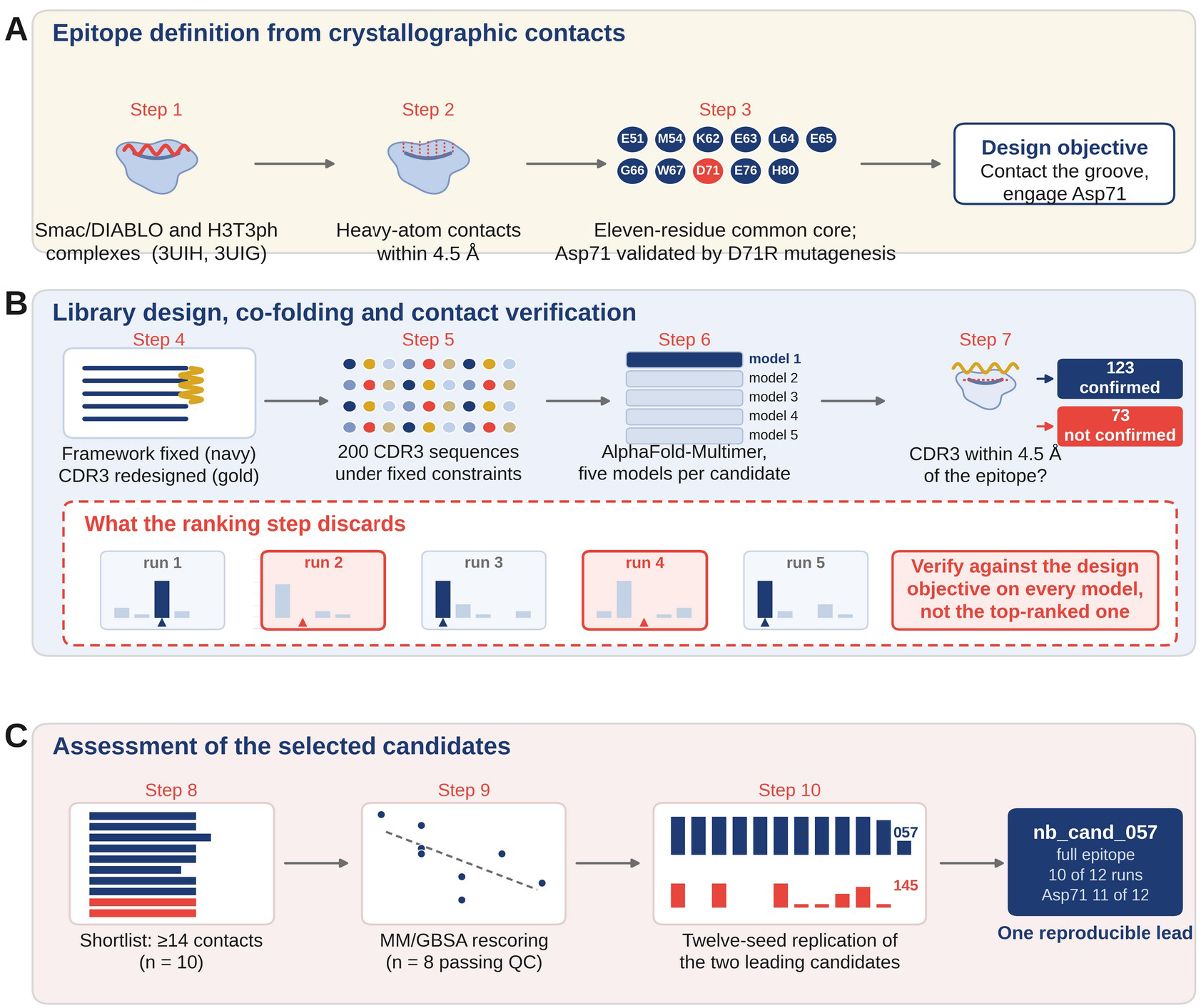
Design and evaluation pipeline. (A) The epitope was defined from crystallographic contacts rather than adopted from the literature: survivin residues within 4.5 Å of the bound peptide in both the Smac/DIABLO (3UIH) and H3T3ph (3UIG) complexes give an eleven-residue common core, of which Asp71 carries independent mutagenesis validation. (B) Two hundred CDR3 loops were designed on a fixed humanised VHH framework and co-folded with the survivin construct; AlphaFold-Multimer produces five models per candidate and the pipeline carries forward the top-ranked one, whose CDR3 is then tested for contact with the epitope. The inset shows what that ranking step discards, measured on the seeded round of 31 sequences, the only set for which all five models were retained: in 9 of 12 apparent failures a model that engaged the epitope had been produced and was rejected. (C) Candidates making fourteen or more epitope contacts were characterised, rescored by MM/GBSA and, for the two leaders, re-predicted under twelve model random seeds, which leaves one reproducible candidate. Step numbers correspond to Sections 2.1 to 2.9.

## 2. Materials and methods

### 2.1 Epitope definition

We selected human survivin (BIRC5; UniProt O15392) as the target, and the BIR-domain groove recognising the free N-terminal alanine of Smac/DIABLO and of the H3T3-phosphorylated histone tail as the epitope. We defined the epitope computationally rather than adopting it from the literature. We retrieved three structures from the RCSB Protein Data Bank: 3UIH (Smac/DIABLO peptide complex), 3UIG (H3T3ph peptide complex) and 1F3H (unliganded survivin). For each liganded structure we recorded every survivin residue with any heavy atom within 4.5 Å of any peptide heavy atom as a contact residue, using Biopython’s NeighborSearch module. Taking the intersection of the two contact sets, that is the residues engaged in both complexes and therefore part of the shared recognition surface, gave an eleven-residue common-core epitope: Glu51, Met54, Lys62, Glu63, Leu64, Glu65, Gly66, Trp67, Asp71, Glu76 and His80.

Two properties of this surface guided the design. It is net acidic, five of eleven residues being aspartate or glutamate, and it presents an aromatic and hydrophobic wall (Trp67, Met54, Leu64) with a single flexible glycine. The D71R substitution abolishes both survivin-Smac binding and anti-apoptotic activity (Song, Yao and Wu, 2003), so we treated Asp71 engagement as a functional criterion throughout. Zinc-coordinating residues were used as landmarks to confirm numbering and not treated as epitope.

### 2.2 Nanobody framework

We used the humanised VHH scaffold h-NbBcII10FGLA, taking its sequence from PDB 3EAK and verifying its identity by pairwise alignment against the parental structure 3DWT, which returned exactly the thirteen substitutions described by Vincke et al. (2009). CDR3 was located programmatically using the conserved cysteine preceding it and the WGQGT motif beginning framework region 4, identifying the donor loop as AAVRGYFMRLPSSHNFRY (18 residues). Every residue outside this window was held fixed across all designs. CDR1, CDR2 and the scaffold are therefore identical in all 200 candidates, and any interface they contribute cannot be credited to the redesign.

### 2.3 Library design

We generated a library of 200 CDR3 sequences by weighted random sampling from an amino-acid pool built to complement the epitope. Arginine and lysine were prioritised for salt bridges to the acidic residues, with glutamine and asparagine as hydrogen-bond donors; tyrosine, phenylalanine, tryptophan, leucine, isoleucine and valine for packing against Trp67, Met54 and Leu64; glycine, serine, alanine and threonine for flexibility near Gly66; and aspartate, glutamate, serine, threonine, asparagine and glutamine as secondary polar contacts to Lys62 and His80. Cysteine was excluded to avoid stray disulfides.

Sequences were accepted only at length 12 to 20 residues, net charge +1 to +5, at most two prolines, at most four repeats of any residue and at most three consecutive hydrophobic residues. Generation used a fixed random seed, so the library regenerates identically.

### 2.4 Complex prediction

We extracted a survivin construct spanning residues 21 to 110 of 1F3H chain A (90 residues) programmatically, defined as the epitope span plus a 30-residue margin on each side, and checked at runtime that all eleven epitope residues fell inside the window. We paired each candidate nanobody with this construct and co-folded the pair with ColabFold 1.6.2 (Mirdita et al., 2022) using AlphaFold-Multimer v3 (Evans et al., 2021), in unpaired alignment mode with three recycles. Five models were generated per complex and re-ranked by the multimer metric, and we carried the top-ranked model forward.

Unpaired mode was used because no coevolutionary relationship exists between a de novo binder and its target. Single-sequence mode was tried first and abandoned: it strips the alignment from survivin as well, and produced unusable predictions (ipTM 0.07 to 0.13).

### 2.5 Epitope contact verification

We did not accept a candidate on interface confidence alone. For every top-ranked model we counted the nanobody residues with any heavy atom within 4.5 Å of any epitope heavy atom, and recorded separately whether any of those contacts came from the redesigned CDR3 loop. A candidate passed only if its CDR3 contacted the intended epitope. A high-confidence interface anywhere else on survivin counted as a failure.

Of the 200 designed candidates we co-folded and assessed 196. Four were lost when a compute session was interrupted mid-batch and are excluded from all analyses. Ten Round 1 candidates predating the contact check (nb_cand_001, 002, 005, 006, 007, 009, 014, 016, 018 and 019) were re-folded to complete the set, and the re-folded values are the ones used throughout. Including a 20-variant mutation and crossover round, reported in Supplementary Section S5, we computed 226 complexes at this stage, before the seeded round and the replication described below.

### 2.6 Interface and model confidence characterisation

We characterised the ten candidates with the highest epitope contact counts in detail, classifying every epitope-residue and nanobody-residue pair within 4.5 Å as a salt bridge, hydrogen bond, hydrophobic or aromatic contact under the distance criteria given in Supplementary Methods. No angular criterion was applied to hydrogen bonds, since the models are unrelaxed and carry no hydrogens; reported hydrogen bonds are distance-defined candidates.

Each contacting nanobody residue was assigned to CDR1, CDR2, CDR3 or scaffold to quantify the fraction of interface contributed by the redesigned loop. Epitope coverage, meaning how many of the eleven epitope residues a candidate engages, was recorded separately from contact count, which counts nanobody residues and can be high for a design concentrated on a few hotspots.

We repeated that partition across every archived predicted complex rather than the shortlist alone, so that the composition of an interface could be related to interface confidence across the library. The same 4.5 Å heavy-atom criterion and the same region boundaries were applied to all 109 archived complexes, counting both the whole nanobody-survivin interface and the part of it falling on the eleven epitope residues. Of those 109, five are Round 2 variants whose loop sequence is not part of the library record and which therefore carry no CDR3 boundary, so the partition is reported for the remaining 104. Only contact-confirmed candidates were archived, so this analysis is conditioned on candidates that made predicted epitope contact; no coordinates survive for the candidates that did not, and interface confidence within the archived set spans 0.15 to 0.69 against 0.15 to 0.78 across the library.

Per-residue model confidence was partitioned by region. AlphaFold writes pLDDT into the B-factor field, so for each complex it was averaged separately over the designed CDR3 and the fixed framework. Since the framework is identical in every candidate, this separates confidence in the borrowed scaffold from confidence in the region the design changed.

To test whether the redesigned loop compromises the scaffold, the ten nanobodies were folded again as isolated chains with no target present, alongside the parent h-NbBcII10FGLA sequence carrying its original CDR3 as a control. All eleven were predicted in one run under identical settings with the ColabFold monomer model and three recycles. The control is necessary because a low CDR3 pLDDT means little on its own: natural CDR loops are mobile and score low, so the question is whether a designed loop scores like a real one. These are monomer predictions from a different network and separate alignments, so they are not on a common scale with the complexes, and candidates are compared only within this run.

### 2.7 Energetic rescoring

We rescored the shortlisted complexes with a single-trajectory MM/GBSA approximation in OpenMM 8.5. The quantity computed is

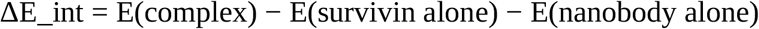

with all three terms evaluated on the same set of minimised complex coordinates. Because the two isolated chains are not re-minimised, every bonded contribution cancels exactly. What survives is the direct interaction between the chains plus the change in solvation free energy on binding.

The solvation term uses the Generalised Born approximation rather than a Poisson-Boltzmann solve, which would have been prohibitive on CPU at this scale. The trade is accuracy in the polar solvation term, examined directly in Section 3.5.

Structures were protonated at pH 7.0 with PDBFixer and built with the amber14 force field and GBn2 implicit solvent, with no nonbonded cutoff and no constraints. Each complex was energy-minimised with all heavy atoms harmonically restrained at 5 kcal mol⁻¹ Å⁻Z, letting steric clashes relax without letting the predicted binding mode drift. Interaction energy was then computed on those minimised coordinates with the restraint removed.

Minimisation length was determined empirically at 500 steps; the convergence series is given in Supplementary Methods.

The survivin receptor is the same molecule in every calculation, so its isolated energy serves as an internal control. The threshold was not set in advance and the reference distribution is that of the runs retained, so this is a post hoc rule. Across the eight successful runs it was −10,689 ± 653 kJ mol⁻¹. Two runs fell far outside that range and were excluded. nb_cand_129 returned +4,290,418 kJ mol⁻¹ (z = +6588) and terminated in 69 s against roughly 280 s for the others, indicating a diverged minimisation; nb_cand_169 returned −2,611 kJ mol⁻¹ (z = +12.4), indicating a strained receptor conformation. Neither failure would have been visible from the final interaction energy alone.

We computed no entropy term and sampled no conformational ensemble, so these are relative, enthalpy-dominated interaction energies rather than binding free energies, and are not converted to dissociation constants.

To establish what the sign of these values means, the calculation was repeated with the Generalised Born force in its own force group, separating the solvation term from the intermolecular gas-phase term. All other settings were unchanged, so this execution also serves as an independent replicate.

### 2.8 Seeded redesign round

We carried out a third design round seeded on a single candidate rather than sampling the library afresh. The seed was nb_cand_145, at that point the only shortlisted candidate returning a negative interaction energy. We fixed the three loop positions the interface analysis identified as functionally important, the two glutamates forming its salt bridges and the single leucine contacting Asp71. We then mutated two or three of the remaining positions per variant, drawing filler positions three times as often as positions already contacting the epitope.

The round carried three arms of ten variants. One added acidic residues, extending the chemistry the seed already used. One added basic residues, the chemistry the library was designed around. One mutated the same positions to neutral polar residues, holding net charge at the seed’s value.

The third arm is the control. Introducing a charged residue both creates a possible ion pair and alters the electrostatics of the whole loop, so comparing a charged arm against the seed alone cannot separate those effects; comparing it against an equally perturbed but charge-neutral arm can. The acidic arm necessarily violates the original net charge constraint and was run with that constraint relaxed. Generation used a fixed random seed, and all 31 sequences were co-folded under the protocol of Section 2.4.

### 2.9 Replication of the leading candidates

One run of this pipeline is five predicted models followed by selection of the top-ranked one. Replication therefore means repeating that whole procedure, not re-reading a single structure. We co-folded nb_cand_145 and nb_cand_057 twelve times each by setting ColabFold’s number of seeds to twelve, which repeats the five-model prediction under twelve model random seeds. Every other setting matched Section 2.4. For each seed we selected the model with the highest 0.8 × ipTM + 0.2 × pTM, the criterion ColabFold uses, and recorded epitope contact for that model and for the best of the five that seed produced. The difference between those two measures what the selection step discards.

The twelve seeds share a single multiple sequence alignment, computed once at the start of the run, whereas the original predictions were made weeks earlier against a different state of the alignment server. This experiment therefore captures variability from the model alone and is a lower bound on the run-to-run variability actually encountered.

We also recorded epitope contact for all five models of every candidate in the seeded redesign round, 155 structures in total, to quantify how often the selection step discards a model that engages the epitope in favour of one that does not.

### 2.10 Statistical analysis and software

Rank correlation is reported as the primary analysis throughout, with Pearson’s *r* given alongside where a linear association is also of interest, since neither epitope contact count nor interface confidence is normally distributed. Correlation coefficients carry 95% confidence intervals from the Fisher *z* transformation, checked against 20,000-sample bootstrap intervals; proportions carry Wilson score intervals; Mann-Whitney comparisons are reported with medians, interquartile ranges and a rank-biserial effect size; and Fisher’s exact tests are reported with the conditional maximum-likelihood odds ratio and its exact confidence interval, which is unbounded above where a cell is zero. Design variables were entered together in a logistic regression of predicted epitope contact on net loop charge, loop length, proline count, aromatic residue count and interface confidence, with batch entered both as a linear term and as indicator terms and tested by likelihood ratio. A different ten Round 1 candidates, whose loop sequences were not carried into the batch ranking files, were recovered from the Round 1 record, so every sequence-based analysis runs on all 196 candidates rather than 186. These are not the ten re-folded in Section 2.5; the two sets do not overlap. Significance is reported at α = 0.05, non-significant results are reported as such, and no correction for multiplicity is applied, so where several measures are compared within one experiment we say so and interpret accordingly. Software: ColabFold 1.6.2 (Mirdita et al., 2022), OpenMM 8.5 and PDBFixer (Eastman et al., 2017), Biopython (Cock et al., 2009), NumPy (Harris et al., 2020), pandas (McKinney, 2010), SciPy (Virtanen et al., 2020), statsmodels (Seabold and Perktold, 2010), Matplotlib (Hunter, 2007) and UCSF ChimeraX (Pettersen et al., 2021). Co-folding ran on NVIDIA T4 GPUs, MM/GBSA on CPU.

## 3. Results

### 3.1 Library evaluation and epitope engagement

Of the 200 designed candidates, we co-folded 196 against the survivin construct and assessed them for epitope contact. Interface confidence spanned ipTM 0.15 to 0.78 across the library, and epitope contact counts spanned 0 to 16 nanobody residues. A total of 123 candidates (62.8%, 95% CI 55.8 to 69.2) made predicted contact with the intended epitope through the redesigned CDR3 loop, verified in the model. The remaining 73 (37.2%, 95% CI 30.8 to 44.2) did not. Seventy-two of those produced predicted complexes in which no nanobody residue came within 4.5 Å of any epitope residue, meaning the model had placed the designed nanobody elsewhere on the survivin surface. The other, nb_cand_163, made a single epitope contact from a region outside the redesigned loop, which the criterion does not credit to the design. Contact rates varied between batches, from 19% to 50% of candidates failing to engage the epitope, with no trend across the order in which we ran them. Nothing in the design or the order of running distinguishes the batches, and the regression below finds no batch effect, so we take the variation as sampling noise; with ten batches of twenty the study could not detect a small systematic one.

One design variable did track engagement. Loops were constrained to net charge +1 to +5, and within that range net charge tracked predicted epitope contact count (Spearman ρ = +0.319, 95% CI +0.19 to +0.44, *P* = 5.3 × 10^−6^, *n* = 196). The miss rate fell across the sampled range, from 47.5% at net +1 (19 of 40) to 8.0% at net +5 (2 of 25), with a single reversal at net +3. The constraint was a hard filter, so every candidate was cationic by construction and this is a relationship within the designed range, not evidence that charge alone suffices.

Entering the design variables together shows that this is not a proxy for loop size or composition. In a logistic regression of predicted epitope contact on net charge, loop length, proline count, aromatic residue count and interface confidence across all 196 candidates, net charge was the only term to reach significance, at an odds ratio of 1.48 per unit charge (95% CI 1.14 to 1.91, *P* = 0.003). Loop length and aromatic content fell just short, in opposite directions (odds ratio 1.14 per residue, 95% CI 0.99 to 1.30; and 0.83 per residue, 95% CI 0.68 to 1.02). Interface confidence contributed nothing once the design variables were held fixed (odds ratio 1.00 per 0.1 ipTM, 95% CI 0.76 to 1.30, *P* = 0.99). Batch had no effect, whether entered as a linear term (*P* = 0.46) or as ten indicator terms (likelihood ratio χ^2^ = 9.98, df = 10, *P* = 0.44), which gives no evidence of a batch effect, although ten batches of twenty is little power to find a small one. The full model is given in Supplementary Table S3. It explains little in absolute terms, McFadden pseudo-*R*^2^ = 0.063 with likelihood ratio *P* = 0.006: the design variables we controlled account for a small part of whether a loop reaches its target.

### 3.2 Interface confidence does not predict epitope engagement

Across all 196 evaluated candidates, interface confidence and predicted epitope contact count were not associated (Spearman ρ = +0.100, 95% CI −0.04 to +0.24, *P* = 0.164; Pearson *r* = +0.130, 95% CI −0.01 to +0.27, *P* = 0.070, *r*^2^ = 0.017). The Pearson coefficient corresponds to 1.7% of the variance in epitope engagement, with an interval that includes none of it (Figure 2A). Against the binary outcome the association is absent altogether: median ipTM is 0.23 (IQR 0.20 to 0.30) among the 123 candidates whose redesigned loop contacted the epitope and 0.22 (IQR 0.20 to 0.28) among the 73 whose did not (Mann-Whitney *P* = 0.759; point-biserial *r* = −0.007, *P* = 0.918).

**Figure 2.**
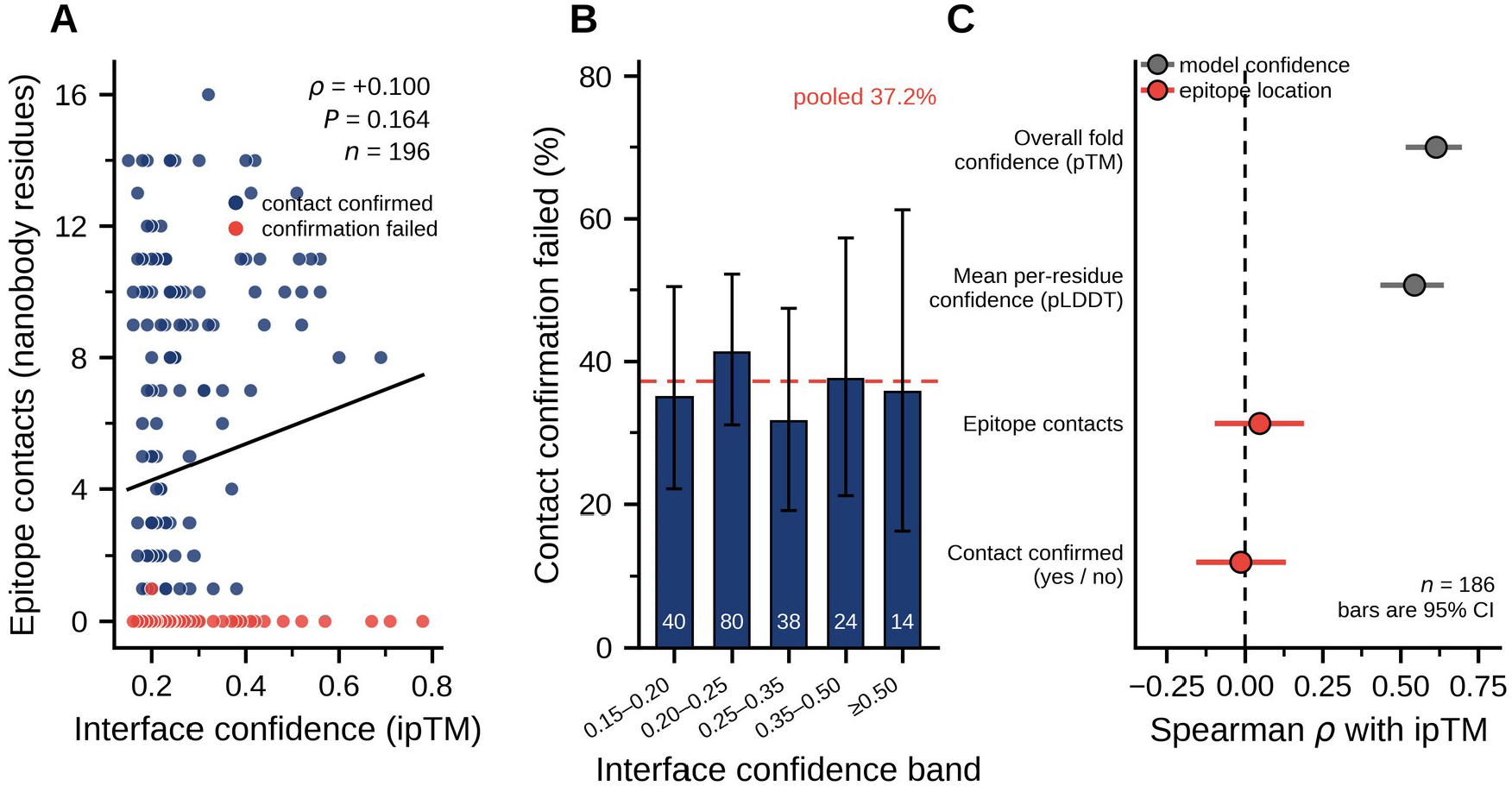
Interface confidence does not predict epitope engagement, and what it does measure is model confidence. (A) Predicted interface confidence (ipTM) against the number of nanobody residues contacting the target epitope, for all 196 evaluated candidates. Navy points denote candidates whose redesigned CDR3 loop was verified in the predicted model to contact the epitope; coral points denote candidates failing that check, whose predicted interface lay elsewhere on the survivin surface. The solid line is the least-squares fit, drawn to show its slope against the scatter rather than to assert a relationship; Spearman ρ = +0.100, *P* = 0.164. (B) Percentage of candidates failing epitope-contact confirmation, partitioned into interface confidence bands, with 95% Wilson score intervals. Sample size is given on each bar; the dashed line marks the pooled failure rate of 37.2%. The failure rate in the highest confidence band (35.7%, 5 of 14) is indistinguishable from that in the lowest (35.0%, 14 of 40). (C) Rank correlation of interface confidence with the two other confidence estimates the same run produces (grey) and with two measures of where the predicted interface sits (coral), the number of epitope contacts and the binary outcome of contact confirmation, for the 186 candidates whose full confidence output was recorded. Bars are 95% confidence intervals from the Fisher *z* transformation; the dashed line marks zero. Interface confidence is a measure of how consistently a complex was built, not of where on the target it was built.

The failure runs in both directions. The three highest-scoring candidates in the library, nb_cand_130 (ipTM 0.78), nb_cand_147 (0.71) and nb_cand_146 (0.67), made no epitope contact at all. Of the fourteen candidates scoring 0.50 or above, five (35.7%) failed contact confirmation. In the other direction, the candidate making the most epitope contacts in the study, nb_cand_197 with 16, scored 0.32, and all ten candidates reaching fourteen or more contacts fell between ipTM 0.15 and 0.42.

Partitioning the library into confidence bands shows a flat failure rate (Figure 2B). Candidates scoring ipTM 0.15 to 0.20 failed contact confirmation 35.0% of the time (14 of 40, 95% CI 22.1 to 50.5); those at 0.50 or above failed 35.7% of the time (5 of 14, 95% CI 16.3 to 61.2). Intermediate bands span 32% to 41%. Taken as quartiles the comparison is the same: the highest quartile failed at 34.7% and the lowest at 35.4%, an odds ratio for failure in the highest quartile relative to the lowest of 0.97 (95% CI 0.41 to 2.26, Fisher’s exact *P* = 1.000). A candidate from the top of the confidence distribution is no more likely to reach the intended site than one from the bottom. AlphaFold-Multimer is not predicting badly here. It is answering a different question: ipTM reports confidence in an interface, not where that interface sits.

One feature of the library bounds how far this generalises. Interface confidence is low across it: the median is 0.23, 49 of the 196 candidates reach 0.30 and only 14 reach 0.50. The relationship is therefore measured mostly inside a band that would ordinarily be read as low-confidence prediction, and a library whose scores spanned a higher range might behave differently. What can be said within this one is that the failure rate does not fall where the score is highest, and that the candidates at 0.50 and above, the only ones a conventional filter would have kept, failed as often as the lowest-scoring forty.

What the score does measure is visible in the same data (Figure 2C). Across the 186 candidates for which the full confidence output was recorded, interface confidence tracks the other confidence estimates closely, at ρ = +0.614 against pTM (95% CI +0.52 to +0.70, *P* = 1.2 × 10^−20^) and ρ = +0.544 against mean pLDDT (95% CI +0.43 to +0.64, *P* = 1.0 × 10^−15^), while tracking predicted epitope contact at ρ = +0.047 (*P* = 0.527). Neither of the other two scores locates the epitope any better: none of the three separates the candidates whose loop contacted the epitope from those whose did not (ipTM *P* = 0.852, pTM *P* = 0.208, mean pLDDT *P* = 0.439). The failure is therefore not specific to ipTM. Confidence in a predicted complex and the location of the predicted interface behave as near-orthogonal quantities, and no rescaling of a confidence score recovers the second from the first.

Partitioning the archived interfaces gives a structural reading of the same point (Supplementary Table S4 and Supplementary Section S7), with the limit that only contact-confirmed candidates were archived, so all 104 archived complexes with a recorded loop sequence are candidates that reached the epitope. Within that set the redesigned loop contributes a median 50.9% of the whole interface (IQR 39.9 to 64.2), and the share it contributes separates the poses reaching the validated anchor from those that do not: median 61.4% (IQR 51.7 to 72.8) among the 54 complexes contacting Asp71 against 40.0% (IQR 34.1 to 47.1) among the 50 that miss it (Mann-Whitney *P* = 1.2 × 10^−8^). Epitope coverage falls as framework contact rises (ρ = −0.571, 95% CI −0.69 to −0.43, *P* = 2.4 × 10^−10^), which is consistent with a complex in which the shared framework does much of the touching being one in which the nanobody has settled across the survivin surface rather than inserted its loop into the groove. Interface confidence does not register that distinction. It falls slightly as framework contact rises (ρ = −0.321, 95% CI −0.48 to −0.14, *P* = 0.0009) rather than being inflated by it, so the shared framework is not quietly raising the score. Both quantities here are read from the same predicted coordinates as the outcome they are compared against, so this describes the geometry of the poses rather than forecasting which candidates will succeed, and it says nothing about the 73 candidates that missed, whose coordinates were not archived.

### 3.3 Replication, and the cost of the selection rule

Losing compute sessions forced us to regenerate structures twice, and both accidents bear on how stable the ranking measures are.

Re-folding the ten Round 1 candidates named in Section 2.5 from identical input several weeks later separated the two measures. Interface confidence held: nine of the ten reproduced to within 0.011 ipTM. The exception, nb_cand_002, moved from 0.196 to 0.370 and rose to the top of its batch while contacting no epitope residue. Epitope contact held less well: nb_cand_038 reproduced its confidence closely, 0.15 to 0.14, while its contact count moved from 14 to 12. Its shortlist admission rests on the original count of fourteen; the re-fold would have placed it below the threshold, and both values are reported in Supplementary Table S2. A reproducible score is not a reproducible pose, and the pose is what the design is about.

Neither measure is stable enough to discriminate finely between candidates. The confidence values held to within 0.011 for nine of the ten but moved by 0.174 for the tenth, and two contacts is within the reproducibility of the contact count itself, so a ranking that separates candidates by margins of that size is reading noise.

Those observations were accidents. To measure the effect deliberately we co-folded the two shortlist leaders, nb_cand_057 and nb_cand_145, twelve times each under twelve model random seeds, selecting the top-ranked model from each run exactly as the pipeline does (**Figure 3**). They behaved completely differently.

**Figure 3.**
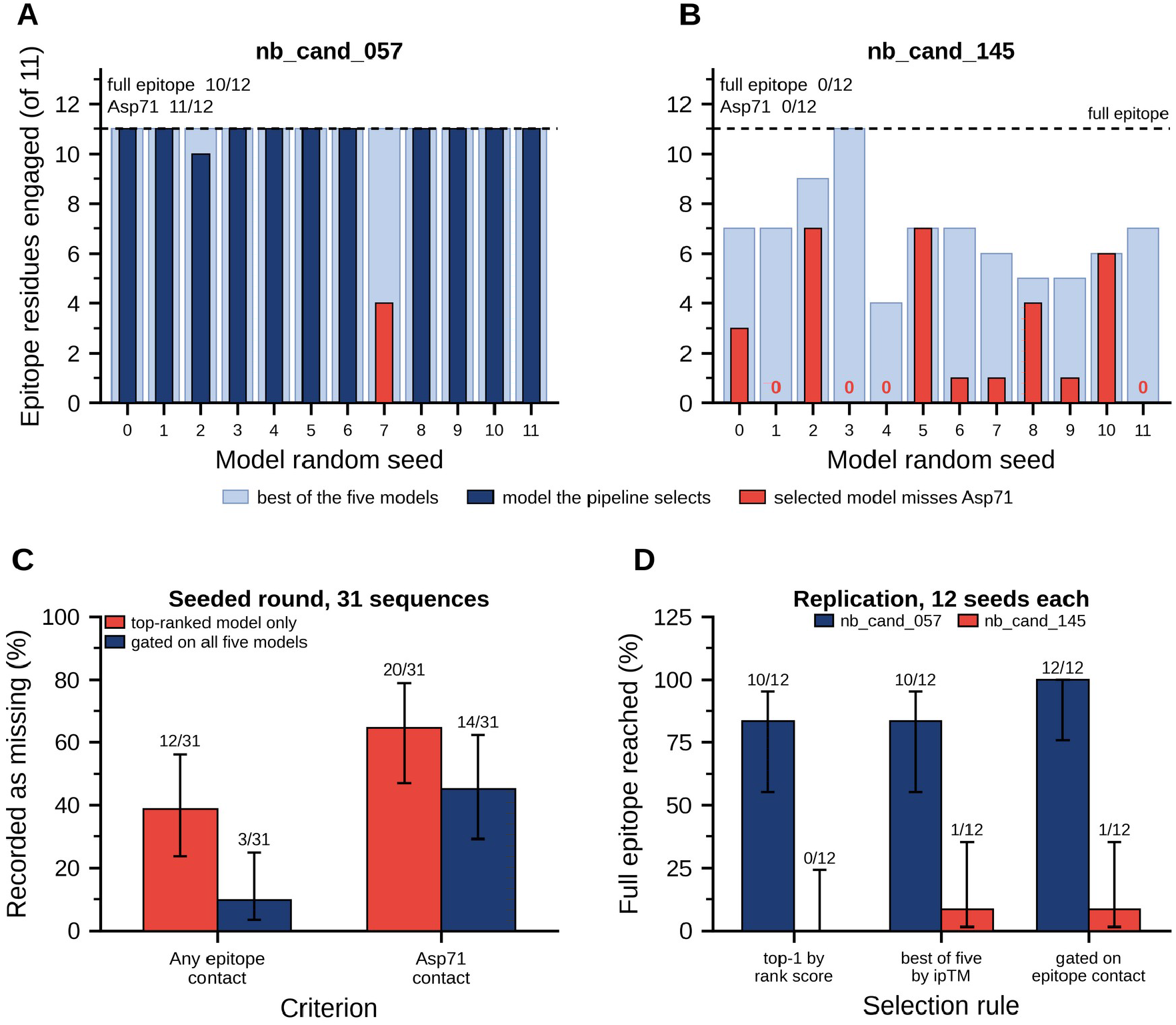
Replication separates two candidates that selection could not, and the selection rule accounts for most of the apparent failures. Each candidate was co-folded twelve times under twelve model random seeds, every other setting held constant. One run of the pipeline is five predicted models followed by selection of the top-ranked one, so each seed is one complete repeat. Dark bars show the epitope residues engaged by the model the pipeline selects; coral marks a selected model that fails to contact Asp71, the epitope residue with independent mutagenesis validation, and a bold zero marks a selected model making no predicted epitope contact at all. Pale bars behind show the best of the five models that seed produced, which the selection step discards. (A) nb_cand_057 reaches the full epitope in 10 of 12 runs and contacts Asp71 in 11 of 12. (B) nb_cand_145 reaches the full epitope in none of the 12 and contacts Asp71 in none, although the best of five reaches the epitope in every run. The two candidates were indistinguishable on the criteria used to select them. (C) In the seeded redesign round, where all five models were retained for all 31 sequences, the proportion of sequences recorded as missing the epitope under the pipeline rule and under a rule that accepts a sequence if any of its five models contacts the epitope, on both the any-contact and the Asp71 criteria. (D) Proportion of the twelve replicate runs reaching the complete eleven-residue epitope under three selection rules. Selecting the best of the five models by interface confidence rather than by the ColabFold rank score changes almost nothing, because the two rules choose the same model in 23 of 24 runs; gating on predicted epitope contact recovers nb_cand_057 completely and does not rescue nb_cand_145. Error bars in C and D are 95% Wilson score intervals and counts are printed above each bar.

nb_cand_057 reproduced. Its selected model made predicted epitope contact in 12 of 12 runs, reached all eleven epitope residues in 10 of 12 (83.3%, 95% CI 55.2 to 95.3) and contacted Asp71 in 11 of 12 (91.7%, 95% CI 64.6 to 98.5), with a median of five interfacial salt bridges (IQR 4 to 5). Its archived result is not one fortunate prediction.

nb_cand_145 did not. Its selected model reached the epitope in 8 of 12 runs but engaged all eleven residues in none and contacted Asp71 in none (both 0 of 12, 95% CI 0.0 to 24.2), with a median of one epitope residue (IQR 0 to 4.5) and zero salt bridges (IQR 0 to 0). Its best selected model across twelve runs reached seven of eleven. The archived structure on which every reported property of this candidate rests, eleven epitope residues, Asp71 contact, three salt bridges and the only negative interaction energy in the study, was never recovered. The gap between the two candidates is large and unambiguous. Epitope residues engaged: median 11.0 (IQR 11 to 11) against 1.0 (IQR 0 to 4.5), Mann-Whitney *P* = 4.3 × 10^−5^, rank-biserial effect size +0.95. Asp71 contact: 11 of 12 against 0 of 12, Fisher’s exact *P* = 9.6 × 10^−6^; the odds ratio has no finite estimate because nb_cand_145 contacted Asp71 in no run, and its exact 95% confidence interval runs from 9.2 upwards.

Nothing in the criteria we applied anticipated this. Both engaged all eleven epitope residues, both contacted Asp71, and they ranked first and second on interaction energy with nb_cand_145 ahead. One property did separate them in the right direction: nb_cand_057 formed five interfacial salt bridges against three, and salt bridge count is the property Section 3.5 finds tracks computed interaction energy. On a single prediction each, that margin is two bonds, and it was not a criterion we applied.

Taking the best rather than the top-ranked of each seed’s five models, both candidates made predicted epitope contact in 12 of 12 runs. The four runs in which the selected model of nb_cand_145 missed the epitope entirely were failures of selection rather than of prediction: a model contacting the epitope existed in every run and was discarded. That is contact of any extent. On the stricter criterion of complete eleven-residue coverage, taking the best of five still leaves nb_cand_145 at 1 of 12, so the two candidates do not converge (Table 1).

**Table 1.** Outcome under three selection rules. The seeded round provides 31 sequences with all five models retained; the replication provides twelve runs of each of two candidates, five models per run, sharing one alignment so the runs are repeats rather than independent samples. Interface confidence was recorded for every model only in the replication, so the ipTM rule cannot be evaluated on the seeded round. The seeded round column counts sequences whose selected model made no predicted contact with any epitope residue; the replication columns count runs reaching all eleven epitope residues, so the two halves of the table use different criteria and are not to be compared across.

| **Selection rule** | **Seeded round:**<br>**sequences missing**<br>**the epitope (of 31)** | **nb_cand_057:**<br>**runs reaching**<br>**the full epitope**<br>**(of 12)** | **nb_cand_145:**<br>**runs reaching**<br>**the full epitope**<br>**(of 12)** |
| --- | --- | --- | --- |
| Top-ranked model only, the rule this pipeline used | 12 (38.7%) | 10 (83%) | 0 (0%) |
| Best of the five models by interface confidence | not recorded | 10 (83%) | 1 (8%) |
| Any of the five models passing epitope-contact gating | 3 (9.7%) | 12 (100%) | 1 (8%) |

The same pattern holds across the seeded redesign round, where we kept all five models for all 31 sequences, 155 structures in total. The top-ranked model missed the epitope for 12 of 31 sequences (38.7%, 95% CI 23.7 to 56.2), close to the 37.2% rate across the library, while only three sequences failed on every one of their five models (9.7%, 95% CI 3.3 to 24.9). Nine of the twelve top-ranked misses had a contacting pose among the five the run had already produced and discarded. On the stricter criterion of Asp71 contact, the top-ranked model missed for 20 of 31 sequences and every model missed for 14 of 31.

Part of that gap is arithmetic. The best of five draws exceeds a single draw whatever is being drawn, so the comparison between the top-ranked model and the best of its five says less than it appears to. The comparison that does not have that defect is between the top-ranked model and the four the ranking rejected, and there the ranking step adds nothing. Top-ranked models engaged a mean of 4.2 epitope residues against 3.6 for the 124 rejected models (Mann-Whitney *P* = 0.610), and missed the epitope at 38.7% against 37.9% (odds ratio 1.03, Fisher’s exact *P* = 1.000). Selecting the top-ranked model is not distinguishable from drawing one of the five at random. That is what the rest of this section rests on: the loss is not that a better model was ranked second, it is that the ranking carries no information about which of the five is right.

Setting the selection rules side by side locates the loss (Table 1, Figure 3C and 3D, and Supplementary Section S6). The replication runs are the only set for which every model’s interface confidence was recorded, so they are the only set that can answer the obvious follow-up: whether re-ranking the five models on ipTM rather than on the ColabFold rank score would have helped. It would not. The two rules select the same model in 23 of 24 runs, and the outcome changes for a single run of nb_cand_145. Two things bound that comparison. The rank score is 0.8 × ipTM + 0.2 × pTM, so the two rules are not independent and close agreement between them is expected rather than surprising; and the 24 runs cover two sequences, so this shows that the two rules coincide in practice, not that no confidence-based rule could do better. What it does establish is that the loss is not an artefact of the particular weighting the pipeline ranks on, and the alternative that does recover the discarded poses is not a score at all.

Gating on predicted epitope contact instead takes nb_cand_057 from 10 of 12 runs to 12 of 12, but leaves nb_cand_145 at 1 of 12, its median epitope coverage rising from one residue (IQR 0 to 4.5) to seven (IQR 5.8 to 7.0) and no further. Contact gating recovers correct poses that ranking discarded; it does not turn an irreproducible candidate into a reproducible one. The two remedies address different failures and neither substitutes for the other: gating fixes the choice made within a run, and replication tests whether the run itself is repeatable.

The highest interface confidence recorded anywhere in the replication, ipTM 0.67 for nb_cand_057 at seed 7, belonged to a model engaging four of eleven epitope residues and missing Asp71. Another model from the same run engaged all eleven. Within a single sequence the score is not useless: across the 60 models of nb_cand_057 it tracks epitope engagement (ρ = +0.482, 95% CI +0.26 to +0.66, *P* = 0.0001), though across the 60 models of nb_cand_145 it does not (ρ = −0.098, 95% CI −0.34 to +0.16, *P* = 0.456).

Neither scalar metric served as a ranking criterion here. Interface confidence does not identify candidates that engage the epitope. Contact count measures the right thing but cannot resolve the top of the distribution, where nine candidates tied at exactly fourteen contacts, a spread smaller than the metric’s own reproducibility. We therefore used verified predicted epitope contact as a binary gate rather than a ranking. Applying it at fourteen or more contacts produced a shortlist of ten library candidates, each admitted on a single prediction, with the consequences set out in Section 3.5.

### 3.4 Interface characterisation of the shortlist

Structural analysis separated candidates that contact count alone had left indistinguishable (Figure 4, Supplementary Table S2). Six engaged all eleven epitope residues, two engaged ten and two engaged nine. The two with the narrowest coverage, nb_cand_129 and nb_cand_173, both failed to contact Asp71. On contact count they were identical to the rest of the shortlist at fourteen each, so excluding them is the first discrimination in this work that rests on functional rather than statistical grounds.

**Figure 4.**
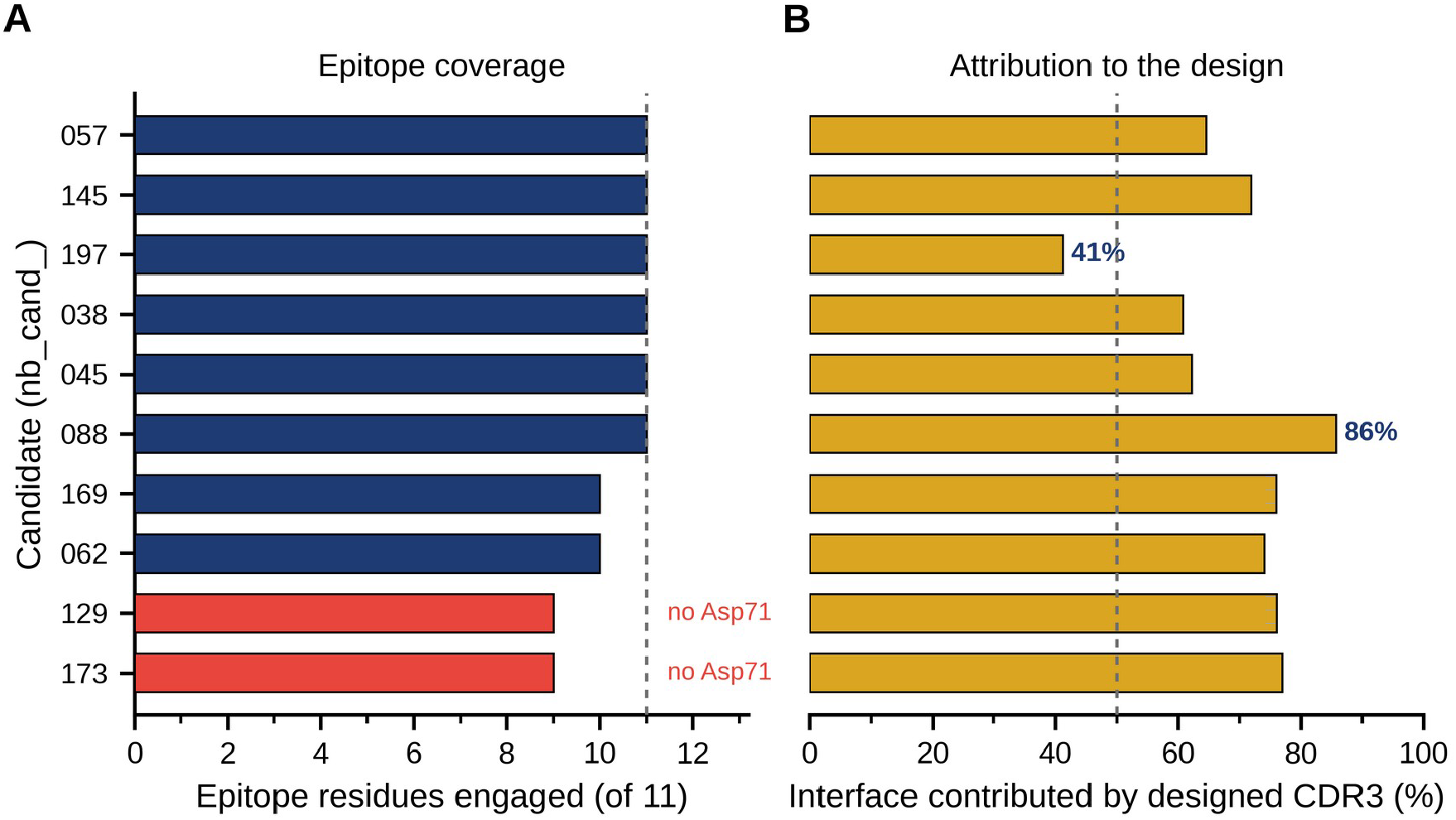
Structural characterisation separates candidates that contact count does not. (A) Number of epitope residues engaged by each shortlisted candidate, of eleven. Coral bars mark the two candidates failing to contact Asp71, the epitope residue with independent mutagenesis validation. (B) Proportion of each interface contributed by the redesigned CDR3 loop; the remainder derives from CDR1, CDR2 and the scaffold, which are identical across all 200 candidates and therefore not attributable to the design. Dotted lines mark full epitope coverage (A) and equal contribution (B). Candidates are ordered by epitope coverage then salt bridge count and labelled by numeric identifier.

Partitioning each interface by region gave a second discrimination. Only CDR3 was redesigned, so interface contributed by other regions is shared by every candidate. The CDR3 proportion ranged from 41.2% to 85.7%. nb_cand_197, which made the most epitope contacts in the study, had the lowest: 20 of its 34 interface residue pairs came from CDR1 and CDR2, so its place at the head of the contact ranking is partly an artefact of counting contacts any candidate on this scaffold could make.

One property held without exception. All 28 salt bridges identified across the ten shortlisted interfaces, which like the hydrogen bonds are distance-defined on unrelaxed models carrying no hydrogens, were formed by CDR3 residues: the designed loop carries the interfacial electrostatics even in candidates where it does not carry the bulk of the contact area. Per-candidate contact counts, epitope residues, salt bridges, hydrogen bonds and CDR3 proportions are given in Supplementary Table S2.

### 3.5 Energetic rescoring and its relationship to interface structure

Among the eight candidates passing quality control, interaction energies ranged from −5.8 to +90.5 kcal mol⁻¹ (Table 2). Only nb_cand_145 returned a negative value.

**Table 2.** MM/GBSA relative interaction energies for the eight shortlisted candidates passing quality control, with structural properties from Supplementary Table S2 for comparison. Candidates are identified by their numeric suffix; all carry the prefix nb_cand_.

| Rank | Candidate | $\Delta E_{\text{int}}$<br>(kcal/mol) | Epitope residues | Asp71 | Salt bridges | CDR3<br>(%) |
| --- | --- | --- | --- | --- | --- | --- |
| 1 | 145 | $-5.8$ | 11 | yes | 3 | 71.9 |
| 2 | 057 | $+13.4$ | 11 | yes | 5 | 64.5 |
| 3 | 197 | $+20.0$ | 11 | yes | 3 | 41.2 |
| 4 | 062 | $+46.5$ | 10 | yes | 2 | 74.1 |
| 5 | 088 | $+49.5$ | 11 | yes | 2 | 85.7 |
| 6 | 045 | $+52.1$ | 11 | yes | 2 | 62.2 |
| 7 | 038 | $+77.7$ | 11 | yes | 2 | 60.7 |
| 8 | 173 | $+90.5$ | 9 | no | 1 | 76.9 |

One structural property tracked the computed interaction energy. Salt bridge count and interaction energy were inversely correlated (Spearman ρ = −0.894, 95% CI −0.98 to −0.51, *P* = 0.003; Figure 5). We treat this as exploratory: it rests on eight candidates drawn from an already-selected shortlist, so the interval around the coefficient is wide and the restricted range means it cannot be extrapolated to the library. Leave-one-out resampling shows it is not the product of a single point, ρ ranging from −0.837 to −0.906 and remaining significant in all eight samples. No other property came close (Supplementary Table S1). It is, however, the largest coefficient from a screen of eight properties on eight candidates, reported without correction for that search, which is a further reason to hold it as exploratory. The relationship is physically sensible: between a cationic loop and an acidic pocket, ion pairing is what offsets the desolvation penalty on binding.

**Figure 5.**
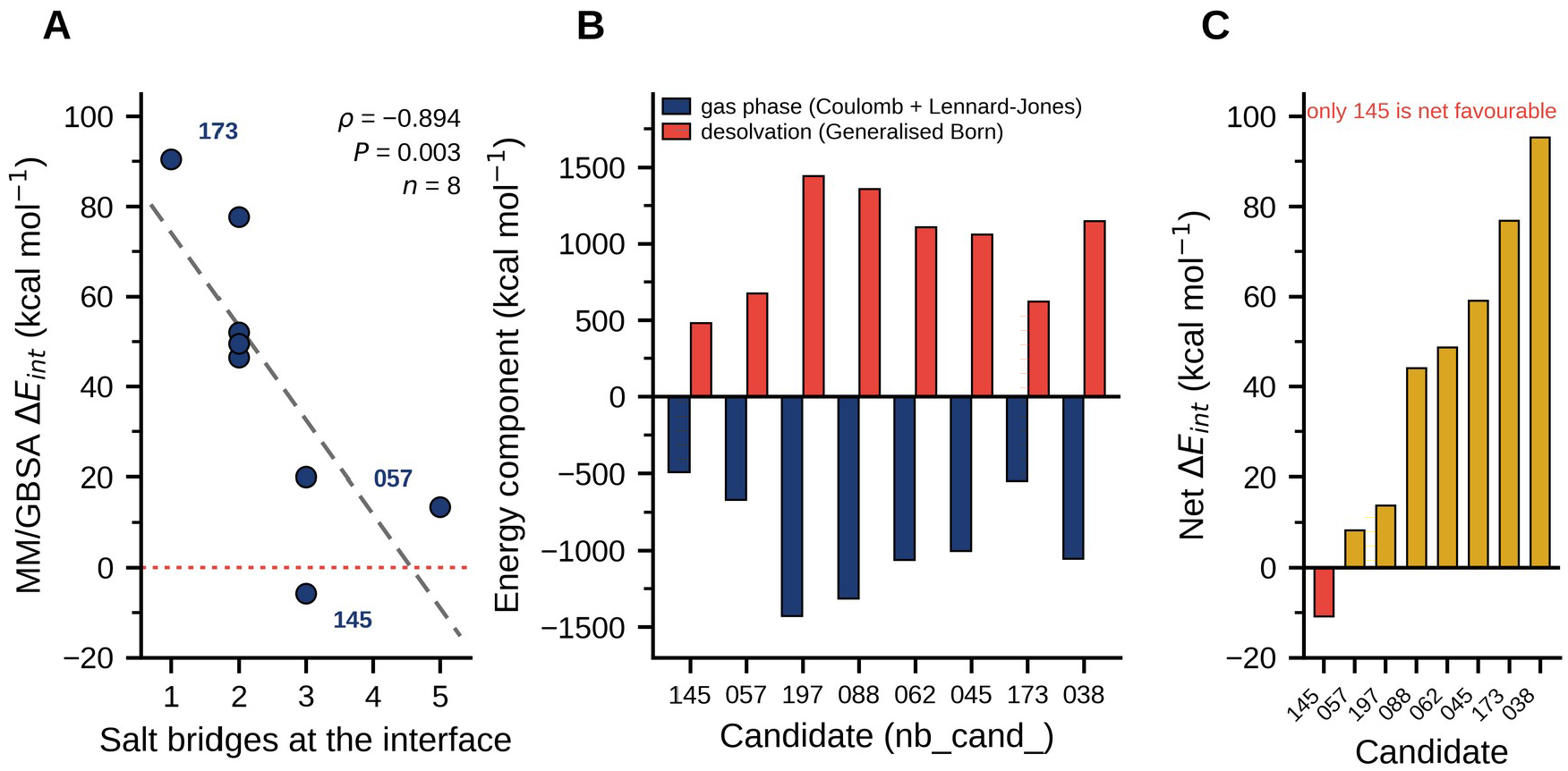
Computed interaction energy tracks salt bridge count and is a small residual of two large opposing terms. (A) MM/GBSA relative interaction energy against the number of interfacial salt bridges, for the eight shortlisted candidates passing quality control. The dashed line is the least-squares fit; the dotted horizontal line marks zero. Spearman ρ = −0.894, 95% CI −0.98 to −0.51, P = 0.003. With eight candidates the association is exploratory. (B) For each candidate, the intermolecular gas-phase term, comprising Coulomb and Lennard-Jones interactions (navy), against the change in Generalised Born solvation free energy on binding (coral). The gas-phase term is favourable for every candidate and the desolvation term unfavourable for every candidate. (C) The net interaction energy, the sum of the two terms in B, on its own scale, which spans roughly a tenth of the range in B. Coral marks the single negative value. Candidates in B and C are ordered by net interaction energy and labelled by numeric identifier. Absolute values are positive for the reasons given in Section 2.7 and only the ordering is interpreted.

A positive interaction energy invites the objection that these complexes are not binding at all, so we decomposed each value into its two terms (**Figure 5B, C**). The intermolecular gas-phase term was favourable for all eight candidates, spanning −492.8 to −1428.2 kcal mol⁻¹. The desolvation term was unfavourable for all eight, spanning +481.9 to +1442.0 kcal mol⁻¹. The reported total is the residual, between 1.0% and 14.1% of the gas-phase magnitude.

A positive total is therefore not evidence of an unfavourable interface. It is the Generalised Born desolvation penalty for burying this many charged groups slightly exceeding the direct electrostatic reward. Since the total is a small difference between two much larger quantities its sign is not robust, and we use these values as a ranking only. Repeating the calculation preserved every sign and the overall rank order (ρ = +0.952, *P* = 0.0003), but moved individual values by 7.8 kcal mol⁻¹ on average and up to 17.6, and exchanged two adjacent pairs; the salt bridge relationship survives that repeat at ρ = −0.830, *P* = 0.011. Net loop charge behaves differently from specific ion pairing, correlating almost perfectly with each component and not at all with their sum (Supplementary Table S1).

When the shortlist was first assessed, nb_cand_145 and nb_cand_057 led on both lines of evidence, ranking first and second on interaction energy and both engaging all eleven epitope residues including Asp71. The replication in Section 3.3 separates them. Every value reported for nb_cand_145 in Table 2 and Supplementary Table S2 comes from a structure twelve further runs did not recover, so it describes one prediction rather than the sequence. That candidate cannot be carried forward, and this study yields one computational lead.

The two candidates excluded on structural grounds were penalised independently on energetic grounds: nb_cand_173 returned the least favourable interaction energy of the eight valid calculations, and nb_cand_129 was the calculation that diverged. One result runs the other way: nb_cand_088 derives the largest proportion of its interface from the designed loop yet ranked fifth of eight on interaction energy.

One caveat covers this section. Every value reported for the shortlist comes from a single predicted structure, and reproducibility has since been tested for two of the ten candidates, one of which failed. Nothing here rules out other shortlisted candidates being as unreproducible as nb_cand_145.

### 3.6 Exploratory forward test of the charge relationship

The salt bridge relationship in Section 3.5 was retrospective, drawn from candidates already selected. We generated a seeded round on nb_cand_145 with three arms of ten variants, adding acidic residues, adding basic residues, or mutating the same positions to neutral residues at constant net charge, and read the charged arms against the neutral control rather than against the seed. This round is an exploratory forward test, not a validation of the salt bridge relationship, for a reason that became clear only afterwards: the seed is itself not reproducible. Section 3.3 shows that the selected model of nb_cand_145 contacts Asp71 in none of twelve repeat runs, so the structure these arms were designed around is a pose the pipeline does not recover. Any result read as a property of that seed is therefore uninterpretable, and we do not read the round that way. What the round can support is a prospective association between net loop charge and predicted epitope engagement, and a second, independent demonstration that the selection step discards correct poses.

Adding basic residues moved predicted epitope engagement in the expected direction; adding acidic residues did not. Contact here means the redesigned loop reaching the epitope, the library criterion, which is stricter than the any-region criterion used for the model counts above. Against the neutral control, the basic arm made predicted epitope contact in 8 of 10 variants against 4 of 10 (odds ratio 5.4, 95% CI 0.60 to 79.9, Fisher’s exact *P* = 0.170), reached a median of 11.0 epitope residues (IQR 3.8 to 11.0) against 1.0 (IQR 0.2 to 4.0) (Mann-Whitney *P* = 0.089), contacted Asp71 in 8 of 10 against 2 of 10 (odds ratio 13.3, 95% CI 1.31 to 240, *P* = 0.023), and formed a median of 2.0 salt bridges against 0.0 (*P* = 0.060). The direction is consistent across all four measures. With ten variants per arm and four measures compared, however, one nominally significant result is weak evidence, and we do not treat the Asp71 comparison as an established finding. The acidic arm matched the control on every measure: epitope contact in 4 of 10 against 4 of 10, Asp71 contact in 1 of 10 against 2 of 10, and medians of zero epitope residues and zero salt bridges in both arms (all *P* ≥ 0.318).

Across all thirty variants net charge tracked predicted epitope engagement (Spearman ρ = +0.535, 95% CI +0.22 to +0.75, *P* = 0.002). This is the arm comparison re-expressed rather than independent evidence, since net charge is what defines the arms, and the range it spans was manufactured by the design of the round. Read that way it is consistent with the relationship found across the library in Section 3.1 and with the original cationic design rule doing what it was intended to do, but it does not test it afresh. The reasoning behind the acidic arm, that extending the seed’s own glutamate-to-Lys62 chemistry would improve it, is not supported.

Re-folded alongside its variants, the seed engaged no epitope residue in the selected model, which is consistent with Section 3.3 and is why the neutral control arm rather than the seed is the only interpretable comparator. The three loop positions held fixed across all three arms were read from that same unrecovered pose, so the scaffold the round was built on is as provisional as the seed itself. A round seeded on nb_cand_057, the candidate that does reproduce, would be the informative version of this experiment and is the natural next computational step.

### 3.7 Fold assessment of the designed nanobodies

Folded without the target, all ten designs returned framework confidence between pLDDT 91.7 and 94.8, against 94.6 for the parent scaffold: a spread of 3.1 units, with no design more than 5 below the parent (Figure 6A). The redesigned loop does not degrade the framework carrying it, and that framework reaches high confidence without the target present.

**Figure 6.**
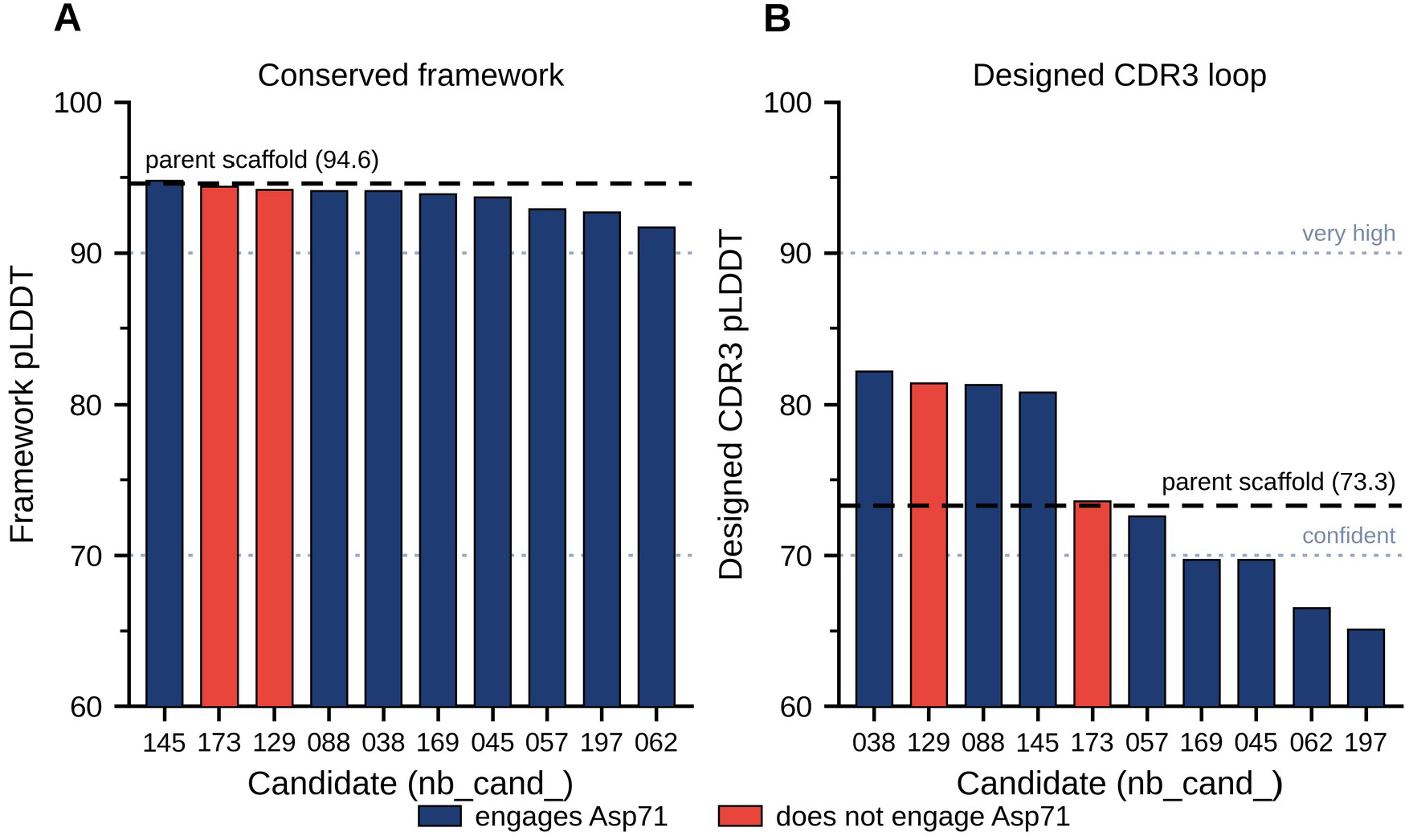
The designed nanobodies fold as well as the parent scaffold without the target. Per-region model confidence for the ten shortlisted designs folded as isolated chains, with the parent h-NbBcII10FGLA scaffold carrying its original CDR3 predicted in the same run as a control (dashed line). (A) Mean pLDDT over the fixed framework. (B) Mean pLDDT over the designed CDR3 loop. Coral bars mark the two candidates excluded for failing to contact Asp71. Dotted lines mark the boundaries of the confidence bands AlphaFold designates very high (90) and confident (70). No design falls more than 5 units below the parent on framework confidence or more than 10 units below it on loop confidence. Candidates are labelled by numeric identifier and ordered by the quantity plotted. Because these are monomer predictions they are not on a common scale with the AlphaFold-Multimer values reported for the complexes, and only comparisons within this figure are interpreted.

The designed loops span pLDDT 65.1 to 82.2 against 73.3 for the parent loop, five above and five below, none more than 10 beneath it (Figure 6B). The parent CDR3 belongs to a nanobody that has been expressed and characterised (Vincke et al., 2009), so the designed loops are no less well defined than a validated loop of the same class under identical conditions. The ranges overlap, which is what a loop predicted as disordered would not do. It does not establish stability, which needs expression and a thermal measurement.

Loop confidence did not identify the candidates meeting the design objective. It bore no relationship to computed interaction energy (ρ = +0.286, *P* = 0.493) or to epitope coverage (ρ = −0.028, *P* = 0.940), and nb_cand_129, excluded for failing to contact Asp71, returned the second-highest loop confidence of any design. Framework and loop confidence correlated within the monomer run (ρ = +0.744, *P* = 0.014).

## 4. Discussion

The practical cost of the confidence result is easiest to see as a cutoff. Filtering at ipTM 0.50 would have selected fourteen candidates, five of which are predicted to bind elsewhere on survivin, and discarded all ten of the candidates making the most epitope contacts. The predicted complexes were not incoherent; they had well-packed interfaces and confidence estimates behaving as the model intends. They were simply confident about something other than location.

This matches the direction of Overath et al. (2025), who found across fifteen targets that ipTM is outperformed by interface-focused alternatives. Their analysis asks whether a design binds at all; ours asks whether it binds in the right place. It is also a caution about how such pipelines get reported. Published workflows filter on interface confidence before testing (Watson et al., 2023; Bennett et al., 2026), and only the candidates that pass are ever characterised, so a filter discarding good binders alongside bad ones would not necessarily be noticed. Evaluating a complete library makes it visible.

Why the score behaves this way is visible in the library itself. Interface confidence is strongly associated with the other confidence estimates the same run produces, at ρ = +0.61 with pTM and ρ = +0.54 with mean pLDDT, and with predicted epitope contact at ρ = +0.05. It is a statement about how consistently a complex was built, and a complex can be built consistently anywhere on the target. None of the three scores separated the candidates that reached the epitope from those that did not, so the problem is not the choice of confidence score but the substitution of confidence for a check against the design objective. That check is available at negligible cost: the epitope is defined before the pipeline runs, and testing a predicted complex against it is a distance calculation on coordinates the run has already written.

The replication result has the sharpest practical consequence. The criteria applied at the time did not separate nb_cand_057 from nb_cand_145, and nb_cand_145 ranked ahead. Only repetition separated them. A candidate selected on one prediction is selected on a sample of one from a distribution nobody measured, and calling it a binder on that basis states more than the evidence carries.

Most apparent failures were failures of selection rather than prediction, and the remedy is nearly free. Checking every model a run produces, rather than the top-ranked one, costs seconds per candidate and would have rescued nine of the twelve failures in the seeded round. The top-ranked model is not a better model: across the seeded round it is indistinguishable from the four the ranking rejected, on epitope residues engaged and on miss rate alike. Re-ranking those same five models on interface confidence does not help either; across the 24 replication runs it selects the model the pipeline already selected in 23 cases, although the rank score is itself mostly ipTM, so the two were never going to disagree often. The general point does not rest on that comparison. A confidence score of any weighting is a statement about the model, and the question the design campaign is asking is about the target.

One qualification. Choosing among models of a single sequence and choosing among candidates in a library are different problems. Within the sixty models of nb_cand_057, interface confidence does track epitope engagement (ρ = +0.482, 95% CI +0.26 to +0.66, *P* = 0.0001), though within those of nb_cand_145 it does not (ρ = −0.098, *P* = 0.456). The score helps a little with the first problem and not at all with the second, so the library-wide result is not a claim that ipTM is meaningless. It is a claim that it does not answer the question a design campaign is asking.

Salt bridge count was the one structural property that tracked computed interaction energy, at ρ = −0.894 and at ρ = −0.830 when the calculation was run again, which tests the numerics rather than any modelling choice, though on eight candidates the association remains exploratory. The decomposition shows why net charge did not. Charge correlates almost perfectly with the favourable electrostatic term and with the desolvation penalty, in opposite directions, so it buys attraction and pays for it almost exactly. Only charge that forms specific pairs survives into the total.

This qualifies the rule that produced the library. Prioritising arginine and lysine was reasoning about a net-acidic pocket, and it worked for what it was reasoning about: net charge predicts whether a candidate reaches the epitope at all, across the library at ρ = +0.319 and prospectively in the seeded round at ρ = +0.535. It does not predict the energy of candidates that get there. The rule was incomplete rather than wrong, and the library alone could not have shown this, because the charge constraint was a hard filter and every candidate was cationic by construction.

Several limitations bound all of this. The largest is that epitope contact is proximity in a predicted model, not a measurement of anything physical. Nothing here was expressed or assayed.

Every structural and energetic value was computed on one predicted structure per candidate, and reproducibility was established for two of the ten shortlisted candidates, one of which failed. The MM/GBSA energies rest on eight candidates from a single minimised snapshot with implicit solvent and no entropy term, and the run-to-run spread of the receptor term alone exceeds the range of the interaction energies it is subtracted from, which is why only the ordering is interpreted and why even that is held as exploratory. The survivin construct is residues 21 to 110 of one chain, predicted as a monomer, so neither the C-terminal dimerisation helix nor the dimer state of the protein is represented. The replication holds one alignment across all twelve seeds, so it captures model stochasticity but not alignment drift. The interface partition in Section 3.2 is bounded more tightly still: only contact-confirmed candidates were archived, so it describes the geometry of complexes that reached the epitope and cannot speak to the ones that did not, and the quantities it relates are read from the same coordinates as the outcome. The scoring relationships were measured for one epitope on one protein, and metric performance varies by target (Overath et al., 2025), so what generalises is the method of measurement rather than any coefficient. Everything reported about these designs was computed; nothing was measured.

Three directions follow.

For design practice, verify against the design objective on every model a run produces, not the top-ranked one, and re-predict any candidate a dozen times before calling it a computational lead. The first uses structures the pipeline already generates; the second costs a few GPU-hours, against the cost of expressing a protein chosen on a sample of one. Where an alternative metric is available, the interface-focused scores evaluated by Overath et al. (2025) merit consideration, though the result here suggests that no score computed from the model alone substitutes for a check against the intended site.

For this target, the immediate experimental step is recombinant expression of nb_cand_057 and a binding measurement against recombinant survivin. A competition assay against a Smac/DIABLO N-terminal peptide would establish whether binding occurs at the intended groove, and alanine substitution of the CDR3 residues forming its predicted salt bridges would test the structural model.

For the method, supplying 3UIH or 3UIG as a template for the survivin chain might bias prediction toward the correct pocket. The effect of that bias on ranking would need separate assessment.

We designed a 200-member nanobody CDR3 library against the survivin BIR-domain Smac/DIABLO groove and evaluated all of it, in order to measure what the criteria used to select designed binders actually select. No association was detectable between interface confidence and whether a design reached the intended site, and none was detectable for the other confidence estimates produced alongside it; what such a score reports is the self-consistency of a predicted complex, not its location. A single run does not establish engagement even when it reports it, and most apparent failures are the ranking step discarding a correct pose the run already produced, a loss that re-ranking on confidence does not recover and epitope-contact gating does. One candidate, nb_cand_057, is predicted to engage the complete epitope including the mutagenesis-validated Asp71 and holds that pose across repeated prediction, while a second that outscored it on every criterion applied at the time does not. nb_cand_057 is a reproducible in silico candidate and nothing more until it is expressed and assayed.

## Data availability

All data, predicted structures and code supporting this study are deposited at Zenodo under doi:10.5281/zenodo.23131327, released under CC BY 4.0 for data and MIT for code. The deposit contains the 196-candidate library sweep, the per-batch pipeline output, the interface characterisation and MM/GBSA rescoring of the ten-candidate shortlist together with its energy decomposition, the twelve-seed replication, the seeded redesign round and the unbound monomer predictions. Predicted complex coordinates were archived only for candidates passing contact confirmation, so the deposit holds 109 structures, covering 104 of the 123 confirmed candidates including all ten on the shortlist and both replication leads, and none of the 73 that failed confirmation. Result tables are plain-text delimited files, provided alongside the consolidated working workbook. All generation and analysis scripts use fixed random seeds and regenerate their inputs identically, the figure pipeline regenerates the six figures reported here from the deposited data, and a verification script re-derives the statistics reported in the Results from the source files and asserts each against the published value.

## Author contributions

Charlotte Gately: conceptualisation, methodology, software, formal analysis, investigation, data curation, visualisation, writing – original draft. Xinyao Guan: conceptualisation, methodology, supervision, writing – review and editing. Xianting Ding: resources, supervision, project administration, writing – review and editing. All authors have read and approved the submitted version and have consented to its deposition as a preprint.

## Funding

This research received no external funding. All structure prediction was carried out on freely available computing resources.

## Conflict of interest

The authors declare no conflict of interest.

## Acknowledgements

We thank our fellow undergraduate researchers in the Institute for Personalized Medicine for their discussion and comments, and the Institute for hosting C.G. during the period in which this work was carried out.

## Notes

https://doi.org/10.5281/zenodo.23131327

### Competing Interest Statement

The authors have declared no competing interest.

