## Supplementary Information for "Benchmarking interface confidence as a selection criterion for computational nanobody design against survivin’s BIR-domain groove"

### Supplementary Methods

#### S1. Contact classification criteria

Every epitope-residue and nanobody-residue pair within 4.5 Å was classified as a salt bridge (charged-group nitrogen to charged-group oxygen within 4.0 Å), a hydrogen bond (nitrogen or oxygen donor-acceptor heavy-atom pair within 3.5 Å), a hydrophobic contact (carbon-carbon within 4.5 Å between apolar residues), or an aromatic contact (ring centroids within 6.0 Å). No angular criterion was applied to hydrogen bonds, since the models are unrelaxed and carry no hydrogens; reported hydrogen bonds are therefore distance-defined candidates.

#### S2. Choice of Generalised Born over Poisson-Boltzmann

The method is MM/GBSA, using the Generalised Born approximation to the solvation term. It is not MM/PBSA, which obtains the same term by solving the Poisson-Boltzmann equation numerically. Generalised Born was used because the calculation had to run on CPU, where a Poisson-Boltzmann solve on a 3,300-atom complex would have been prohibitive at this scale. The trade is accuracy in the polar solvation term, examined directly in Section 3.5 of the main text.

#### S3. Minimisation convergence

Minimisation length was determined empirically. On unminimised models the interaction energy is dominated by clashing atoms and returns values of order $10^7$ kcal mol$^{-1}$. A convergence series on a single candidate gave +52.78, +45.55, +43.34 and +41.36 kcal mol$^{-1}$ at 100, 250, 500 and 1000 steps respectively. We adopted 500 steps, since doubling to 1000 changes the result by less than 2 kcal mol$^{-1}$.

### Supplementary Results

#### S4. Per-region model confidence across the shortlist

Across the ten shortlisted complexes the framework averaged pLDDT 80.8 to 87.3, the band AlphaFold calls confident, as expected for a scaffold taken from a deposited structure. The designed loops averaged 45.9 to 62.1, the band it calls low, and the least confident residue in each loop fell between 32.3 and 49.8. The predictor is least certain about the one region the design changed.

That is expected for a de novo loop, which has no alignment support and would be genuinely mobile in a folded antibody. It is not a measurement of instability, but it does bound how firmly the predicted binding mode can be asserted. Loop confidence did not track the selection criteria, showing no relationship with salt bridge count (Spearman $\rho = +0.113$, $P = 0.755$) and only a non-significant trend with computed interaction energy ($\rho = -0.619$, $P = 0.102$). The two leads carried the highest and third-highest loop confidence in the shortlist, nb_cand_145 at 62.1 and nb_cand_057 at 58.7.

### S5. Mutation and crossover round

The mutation and crossover round produced twenty variants from the five leading Round 1 candidates. All ten mutation-derived variants retained verified epitope contact, against eight of ten crossover-derived variants, with both losses being crossovers. The sample is too small for a statistical claim, but the direction fits the expectation that small perturbations of a validated design are more conservative than recombining two designs. No Round 2 variant entered the final shortlist, which was drawn from the library sweep.

### S6. Which model each selection rule chooses

Two sets in this study retained every model a run produced rather than only the top-ranked one, and they answer different parts of the same question. The seeded redesign round retained all five models for all 31 sequences, 155 predictions, but recorded only each model's ColabFold rank, not its interface confidence. The twelve-seed replication retained all five models of each of 24 runs, 120 predictions, with ipTM and pTM for every model, and is therefore the only set in which a confidence-based re-ranking can be simulated. ColabFold orders models by $0.8 \times \text{ipTM} + 0.2 \times \text{pTM}$, so ranking on ipTM alone is not independent of the pipeline rule; the question is how far the two diverge in practice. They select the same model in 23 of the 24 runs, and the one divergence changes the outcome for a single run of nb_cand_145.

The gate that does recover discarded poses is a check against the epitope itself. In the seeded round it reduces the proportion of sequences recorded as missing the epitope from 12 of 31 to 3 of 31, and on the Asp71 criterion from 20 of 31 to 14 of 31. In the replication it takes nb_cand_057 from 10 of 12 runs reaching the complete epitope to 12 of 12, and leaves nb_cand_145 at 1 of 12. The two interventions are not interchangeable: gating corrects the choice made within a run, replication tests whether the run is repeatable at all.

### S7. Composition of the predicted interfaces

Every archived predicted complex was partitioned by which region of the nanobody contributes each interfacial residue pair, under the same 4.5 Å heavy-atom criterion and the same CDR1, CDR2, CDR3 and scaffold boundaries used for the shortlist in Section 2.6 of the main text. Table S4 relates interface confidence to that partition. Only contact-confirmed candidates were archived, so all 104 complexes with a recorded loop sequence are candidates that reached the epitope, interface confidence within the set spans 0.15 to 0.69 against 0.15 to 0.78 across the library, and nothing in this table describes the candidates that missed.

The pattern is consistent across the rows. Interface confidence is unrelated to how much of the interface the redesigned loop contributes in absolute terms, falls as contact from the shared framework rises, and rises with the proportion of the interface the designed loop carries and with the number of epitope residues engaged. A predicted complex in which the framework does much of the touching is one in which the nanobody has settled across the surface rather than inserted its loop into the groove, and such complexes are built with lower confidence overall, which is why the association with framework contact is negative rather than positive.

### Supplementary Tables

**Table S1.** Spearman correlations against MM/GBSA interaction energy across the eight shortlisted candidates passing quality control, and correlations of net loop charge with the two energy components. Salt bridge count is the only structural property with a significant relationship to the net energy.

| Property | ρ | P |
|---|---|---|
| Epitope residues engaged | −0.436 | 0.280 |
| Contact count | −0.436 | 0.280 |
| Hydrogen-bond pairs | −0.024 | 0.954 |
| Proportion of interface from CDR3 | +0.143 | 0.736 |
| Salt bridge count | −0.894 | 0.003 |
| Net loop charge vs gas-phase term | −0.933 | 0.001 |
| Net loop charge vs desolvation term | +0.933 | 0.001 |
| Net loop charge vs net interaction energy | +0.258 | 0.538 |

**Table S2.** Interface characterisation of the ten shortlisted candidates, ordered by epitope coverage then salt bridge count. Candidates are identified by their numeric suffix; all carry the prefix nb_cand_. Contact counts are nanobody residues within 4.5 Å of any epitope residue. The value marked with an asterisk derives from a re-folded structure.

| Candidate | Contacts | Epitope residues | Asp71 | Salt bridges | H-bonds | CDR3 (%) |
|---|---|---|---|---|---|---|
| 057 | 14 | 11 | yes | 5 | 11 | 64.5 |
| 145 | 14 | 11 | yes | 3 | 10 | 71.9 |
| 197 | 16 | 11 | yes | 3 | 10 | 41.2 |
| 088 | 14 | 11 | yes | 2 | 10 | 85.7 |
| 045 | 14 | 11 | yes | 2 | 17 | 62.2 |
| 038 | 12* | 11 | yes | 2 | 13 | 60.7 |
| 169 | 14 | 10 | yes | 4 | 12 | 75.9 |
| 062 | 14 | 10 | yes | 2 | 8 | 74.1 |
| 129 | 14 | 9 | no | 4 | 6 | 76.0 |
| 173 | 14 | 9 | no | 1 | 6 | 76.9 |

**Table S3.** Logistic regression of predicted epitope contact on the design variables, across all 196 evaluated candidates. Net loop charge is the only term reaching significance; interface confidence contributes nothing once the design variables are held fixed. McFadden pseudo-$R^2$ = 0.063, likelihood-ratio P = 0.006. Batch entered as ten indicator terms does not improve the fit (likelihood ratio $\chi^2$ = 9.98, df = 10, P = 0.442).

| Term | Coefficient | Odds ratio | 95% CI | P |
|---|---|---|---|---|
| Net loop charge, per unit | +0.389 | 1.48 | 1.14 to 1.91 | 0.003 |
| Loop length, per residue | +0.127 | 1.14 | 0.99 to 1.30 | 0.062 |
| Proline count, per residue | −0.013 | 0.99 | 0.64 to 1.52 | 0.955 |
| Aromatic count, per residue | −0.190 | 0.83 | 0.67 to 1.01 | 0.069 |
| Interface confidence, per 0.1 ipTM | −0.002 | 1.00 | 0.76 to 1.30 | 0.986 |

**Table S4.** Rank correlation of interface confidence with the composition of the predicted interface, across the 104 archived complexes carrying a recorded loop sequence. Confidence intervals are from the Fisher z transformation.

| Quantity | ρ with ipTM | 95% CI | P |
|---|---|---|---|
| Total interfacial residue pairs | −0.336 | −0.50 to −0.15 | 0.00049 |
| Pairs contributed by the shared framework | −0.321 | −0.48 to −0.14 | 0.00089 |
| Pairs contributed by CDR1 | +0.027 | −0.17 to +0.22 | 0.784 |
| Pairs contributed by CDR2 | −0.311 | −0.48 to −0.13 | 0.001 |
| Pairs contributed by the scaffold | −0.274 | −0.44 to −0.09 | 0.005 |
| Pairs contributed by the redesigned CDR3 | −0.061 | −0.25 to +0.13 | 0.537 |
| CDR3 share of the whole interface (%) | +0.216 | +0.02 to +0.39 | 0.028 |
| Pairs falling on the eleven epitope residues | +0.227 | +0.04 to +0.40 | 0.021 |
| Epitope residues engaged | +0.439 | +0.27 to +0.58 | $3.2 \times 10^{-6}$ |

*All underlying data, structures and analysis code are deposited at Zenodo under doi:10.5281/zenodo.23131327; see Data availability in the main text.*
